# Inferring functionally relevant molecular tissue substructures by agglomerative clustering of digitized spatial transcriptomes

**DOI:** 10.1101/2020.11.09.374660

**Authors:** Julien Moehlin, Bastien Mollet, Bruno Maria Colombo, Marco Antonio Mendoza-Parra

**Affiliations:** UMR 8030 Génomique Métabolique, Genoscope, Institut François Jacob, CEA, CNRS, University of Evry-val-d’Essonne, University Paris-Saclay, 91057 Évry, France

**Keywords:** Spatial Transcriptomics, image analysis, gene expression elements, systems biology

## Abstract

Developments on spatial transcriptomics (ST) are providing means to interrogate organ/tissue architecture from the angle of the gene programs defining their molecular complexity. However, computational methods to analyze ST data under-exploits the spatial signature retrieved within the maps. Inspired by contextual pixel classification strategies applied to image analysis, we have developed MULTILAYER, allowing to stratify ST maps into functionally-relevant molecular substructures. For it, MULTILAYER applies agglomerative clustering strategies within contiguous locally-defined transcriptomes (herein defined as gene expression elements or Gexels), combined with community detection methods for graph partitioning.

MULTILAYER has been evaluated over multiple public ST data, including developmental tissues but also tumor biopsies. Its performance has been challenged for the processing of high-resolution ST maps and it has been used for an enhanced comparison of multiple public tissue biopsies issued from a cancerous prostate.

MULTILAYER provides a digital perspective for the analysis of spatially-resolved transcriptomes and anticipates the application of contextual gexel classification strategies for developing self-supervised molecular diagnostics solutions.

Overall, the development of MULTILAYER anticipates the application of contextual gexel classification strategies for developing self-supervised molecular diagnostics solutions.

## Introduction

Studying complex living systems by the evaluation of the various gene programs defining organ/tissue architecture, is part of the current challenge on Systems Biology. In fact, while till recently accessing at the gene programs in tissues was performed by global (bulk) gene expression analyses, recent advances on single cell transcriptomics managed to move from an “average view” towards a single-cell gene program readouts^1^. This being said, cell dissociation by enzymatic methods - necessary for single-cell assays - tend to modify the transcriptional patterns^2^, it destroys for at least a fraction of the cells composing the tissue, and it does not conserve tissue architecture.

Recent developments on “spatial transcriptomics” (ST)^3^, allows to circumvent the aforementioned technical issues related to single-cell assays, and notably the capacity to conserve the spatial architecture, essential for heterogeneous tissue analysis. This strategy, based on the use of a physical support (DNA array) for capturing local gene expression signatures (mRNA transcriptome) from tissue sections, behaves like a digital camera, allowing to obtain a “digital” view of the molecular programs composing the tissue.

While several computational solutions are currently available for processing ST^4,5^, their analytical pipelines tend to reuse strategies applied for single-cell transcriptomics analyses, namely to consider each of the captured local transcriptomes as independent units during their comparison. Herein we describe MULTILAYER, a stand-alone package allowing to process ST readouts by pattern recognition within contiguous local transcriptomes. Such captured local transcriptomes are defined herein as gexels (gene expression elements), in analogy to pixels commonly described as units composing raster images in digital imaging. Hence, MULTILAYER process ST maps as a digital image on which gexel patterns revealed from agglomerative clustering allows to highlight biologically relevant tissue substructures. MULTILAYER is available through https://github.com/SysFate/MULTILAYER.

## Results

### Normalization, differential Gene expression, co-expression patterns detection and digital tissue partitioning of ST data performed by MULTILAYER

MULTILAYER receives as input, ST matrices composed by spatial coordinates and read counts per genes. These matrices are converted into a grid view, on which each spatial coordinate is associated to a gene expression element (or gexel), composed by read-counts per gene for the local transcriptome. Raw ST maps present variable total read counts per gexel, potentially due to technical concerns during sample preparation (e.g. uneven tissue permeabilization, mRNA capturing, etc). To address this problem, MULTILAYER applies a quantile normalization^6^ across Gexels, generating a uniform total read counts map over the whole grid (**Figure 1**). Normalization performance has been validated by evaluating housekeeping gene expression levels within spatial transcriptome maps and relative to other alternative strategies, like SpatialDE^7^ (**Supplementary Figure 1**).

**Figure 1.**
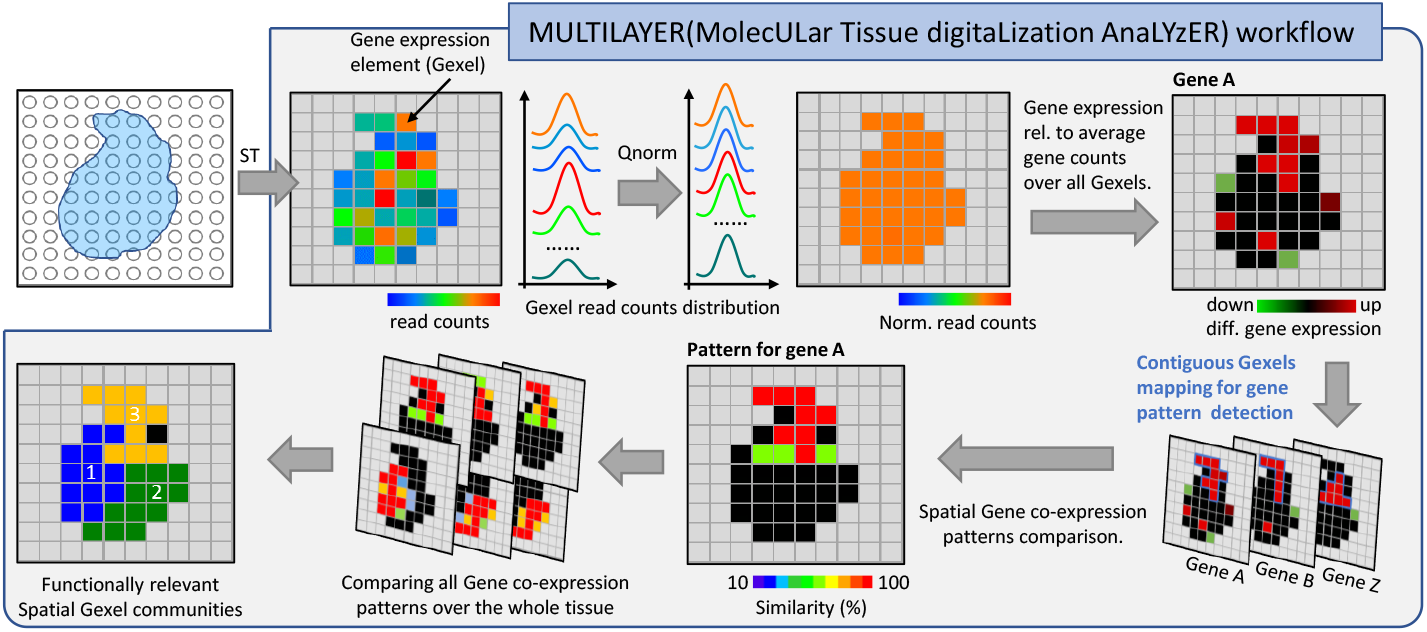
The Molecular Tissue Digitalization Analyzer (MULTILAYER) workflow. MULTILAYER requires as input spatial transcriptomics matrices composed by spatial coordinates harboring read-counts per genes. Each of such coordinates are defined as gexels (gene expression elements) in analogy to pixels composing digital images. At first, MULTILAYER corrects for differences on total read-counts per gexel since such variations are considered as artifactual. Normalized matrices are used for computing differential gene expression relative to the average expression over the whole tissue. Similar to digital image processing, an agglomerative strategy is applied to reveal gene patterns defined by contiguous gexels, which are then compared to reveal spatial gene co-expression patterns, expected to host functionally relevant information. A global comparison of all gene coexpression patterns leads to the partition of the initial spatial transcriptomics map into functionally relevant spatial community regions.

Under the hypothesis that the digital tissue map under study is not homogeneous, we aimed at inferring changes on gene expression in a spatial context. For it, MULTILAYER computes gene expression levels per gexel relative to the gene expression average behavior within the tissue. In analogy to the terminology used on “bulk” gene expression analysis, we describe herein regions with higher levels as up-regulated, or down-regulated when the normalized read counts per gene were above or below the stated average behavior (**Figure 1**). While this analysis is performed per Gexel, MULTILAYER ranks differentially expressed genes on the grounds of the observed number of Gexels, hence providing the user with a quick view of the genes that are over-represented on the digital map on the basis of the relative expression behavior (**Supplementary Figure 2**).

Similar to contextual classification strategies used for image analysis from pixels information^8^, MULTILAYER detects gene expression patterns by the use of an agglomerative strategy over contiguous Gexels. This module generates a cleaner view of over-expressed genes within the tissue. Furthermore, it allows to identify multiple patterns for the same overexpressed gene within the same tissue, which per see is lost in all other strategies relying on aggregating independent Gexels by applying for instance the t-distributed stochastic neighbor embedding (tSNE) method (**Supplementary Figure 3**).

Having detected patterns for all over-expressed genes, MULTILAYER compares their spatial localization to infer their degree of co-expression behavior (Tanimoto and Dice similarity coefficient implemented; see methods on line). By expanding this analysis through the ensemble of over-expressed genes, MULTILAYER generates a graph structure on which nodes corresponds to over-expressed genes within the tissue and edges their similarity coefficient readouts reflecting their degree of spatial co-expression (**Supplementary Figure 4**). By applying the Louvain methodology for community detection^9^, MULTILAYER partitions the digital tissue map into functionally relevant spatial Gexel substructures (**Figure 1**).

### MULTILAYER efficiently partitions digitized tissue maps into relevant functional substructures

To assess the performance of MULTILAYER, we have processed public ST maps issued from a variety of tissues, including human developmental heart samples^10^ and pancreatic tumors^11^. As part of the heart study, MULTILAYER has been instrumental to decorticate the tissue complexity within 19 digital maps covering three described developmental stages (4.5 and 6.5 and 9 post-conception weeks (PCW); **Supplementary Figure 5**). To highlight the performance of MULTILAYER on tissues presenting a high complexity, we have focused our attention on a tissue section collected at nine PCW (Section 15; **Supplementary Figure 5**). After applying quantile normalization on the raw read counts (**Figure 2A**), MULTILAYER detected several over-expressed genes within the heart section, including those coding for the smooth muscle actin ACTA2; one of the components of the elastic fibers (Elastin; ELN), the large abundant protein composing the striated muscle Titin(TTN) or the natriuretic peptide NPPA (**Figure 2B**). The spatial gene over-expression signature for ACTA2 appeared concomitant with that of ELN and distinct to those observed for TTN and NPPA. This observation is confirmed by the spatial gene co-expression analysis performed by MULTILAYER, demonstrating that ACTA2 and ELN present a similarity index > 30% (Tanimoto distance) (**Figure 2C**), also observed for other factors like PXDN, BGN, S100A11, HTRA1, EMILIN1, CXCL12, MYH10 or TAGLN; some of them previously described as being expressed at the heart valve^12–14^. In a similar manner, TTN presented a co-expression signature with NPPA, as well as several other factors like NDUFA4, FHL2, CTNNA1, known to present a specific left ventricle over-expression (as documented on the Genotype-tissue expression portal ^15^).

**Figure 2.**
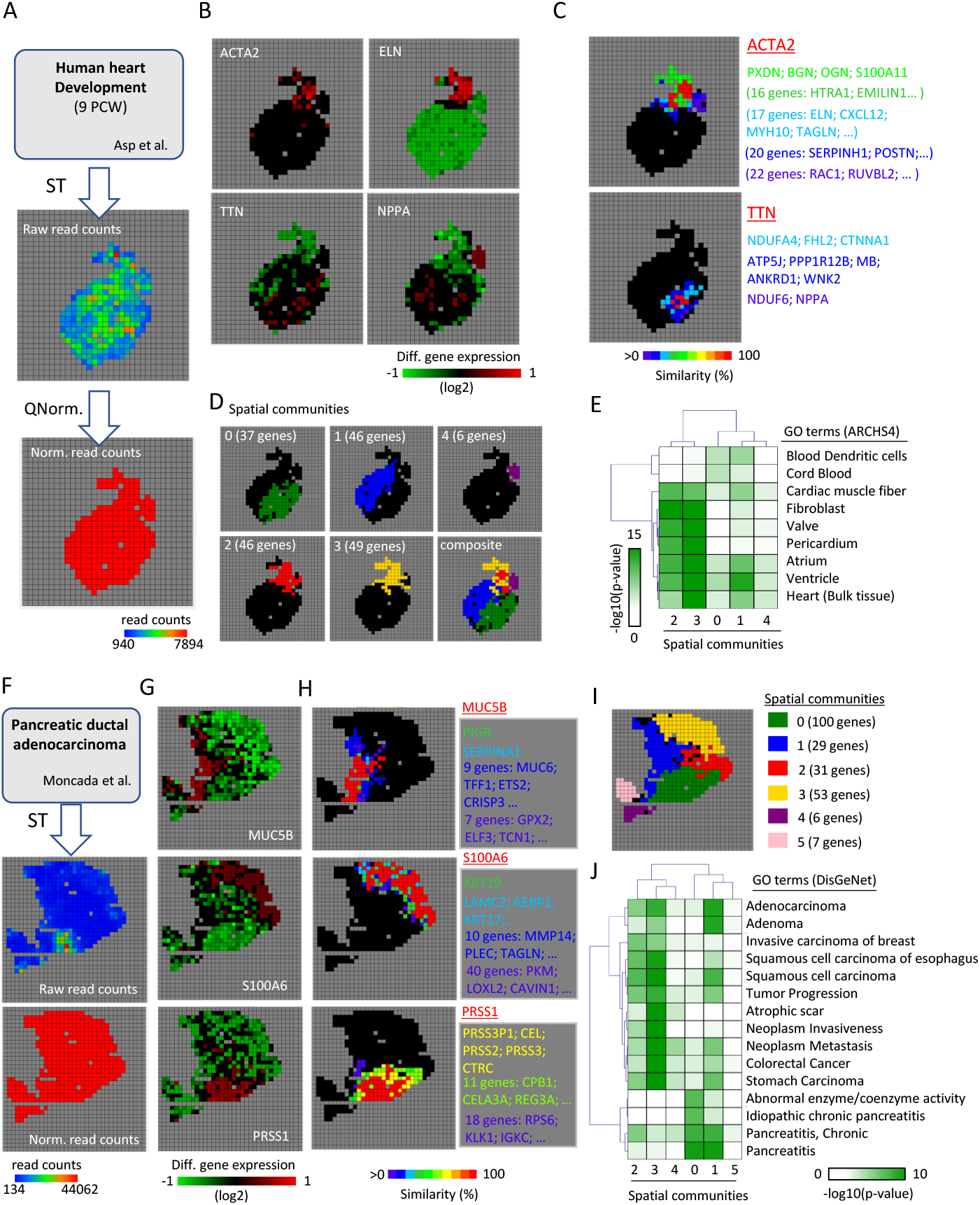
Inferring Functionally relevant tissue substructures with MULTILAYER. **(A)** MULTILAYER normalization of spatial transcriptomics data issued from developing human heart tissue (9 post-conception weeks; M. Asp et al.). **(B)** Differential gene expression signature (relative to the average behavior within the tissue) for the genes ACTA2, ELN, TTN and NPPA as revealed after normalization. **(C)** Spatial gene co-expression analysis performed by MULTILAYER for ACTA2 and TTN. Gexels colored in red corresponds to the location of the target genes (ACTA2 or TTN); while the other colored gexels reveals their coexpression pattern (Tanimoto similarity index). **(D)** Spatial community tissue stratification issued from a gene co-expression analysis performed over the whole tissue and all overexpressed genes. The developing human heart tissue map has been stratified on 5 spatial communities (from “0” to “4”), from which two of them present a highly redundant spatial localization pattern (“2” & “3”). **(E)** Gene ontology analysis (ARCSH4 tissue database) for each of the spatial communities retrieved within the developing human heart tissue, as performed by MULTILAYER. **(F)** MULTILAYER normalization of spatial transcriptomics data issued from pancreatic ductal adenocarcinoma tissue (Moncada et al.). **(G)** Differential gene expression signature for the genes MUC5B, S100A6 and PRSS1 after tissue normalization. **(H)** Spatial gene co-expression analysis performed by MULTILAYER for MUC5B, S100A6 and PRSS1. Gexels colored in red corresponds to the location of the target genes; while the other colored gexels reveals their co-expression pattern (Tanimoto similarity index). **(I)** Composite view of the 6 spatial communities detected on the pancreatic adenocarcinoma tissue from a gene co-expression analysis performed over the whole tissue and all overexpressed genes. **(J)** Gene ontology analysis (DisGeNET database) for each of the spatial communities displayed in (I), as performed by MULTILAYER. ARCHS4 tissues: GO terms database issued from Massive Mining of Publicly Available RNA-seq Data from Human and Mouse (*Lachmann A. et al; Nat. Comm. 2018*). DisGeNET: Disease-Gene association discovery platform (Piñero J. el al; NAR 2020).

By extending the gene co-expression pattern detection over the whole tissue and applying spatial communities partitioning, MULTILAYER revealed the presence of five communities, which can be summarized within 3 major distinct tissue substructures (**Figure 2D**). Gexel communities “0” and “1” corresponds to two distinct regions, associated to the left and right ventricle and atrium of the heart (**Figure 2E**). In contrast, communities “2” and “3” present a redundant spatial location (**Figure 2D**), which were functionally related to pericardial tissue but also to the heart valve (**Figure 2E**).

To further illustrate the performance of MULTILAYER on other type of tissues, we have analyzed Pancreatic ductal adenocarcinoma ST maps^11^. For it, we have first normalized the publicly available ST map (**Figure 2F**), then MULTILAYER detected a variety of spatially over-expressed genes, among them the mucin family member MUC5B, the S100 Calcium Binding Protein A6 (S100A6) or the serine protease PRSS1 (**Figure 2G**). Notably, these three over-expressed genes present a completely distinct spatial behavior, further confirmed by their gene co-expression patterns as inferred by MULTILAYER (**Figure 2H**). Such distinct spatial pattern behavior is in agreement with their previously described functional role, since MUC5B has been over-expressed on pancreatic duct^16^, S100A6 has been associated to pancreatic cancer development^17^ and PRSS1 is expressed on normal pancreas tissue since it codes for trypsinogen, the enzyme secreted by this organ. Beyond these three distinct regions, MULTILAYER inferred up to 6 gexel communities (**Figure 2I**), which can be summarized on 4 functionally relevant regions. Gexel community “0”, has been associated to functional terms like Pancreatitis, or abnormal enzyme activity, most likely due to the fact that mutation on PRSS1 were associated with hereditary pancreatic disorders^18^. This spatial region has been characterized as “normal pancreas tissue” as part of the histological annotation described by Moncada et al^11^, which is in agreement with the MULTILAYER functional annotation, devoid of tumor-related terms (**Figure 2J**). In contrary, gexel community “3” has been strongly associated to disease terms like “Adenocarcinoma”, “tumor progression”, or “Neoplasm metastasis”; in agreement with the histological annotation described by Moncada et al^11^. Similarly, gexel communities “2” and “1” were associated to “cancer related terms”, but with a lower confidence, in agreement with the aforementioned histological differences (Gexel community “1” described as duct epithelium, and “2” as Stroma^11^).

Overall, MULTILAYER allowed to perform an automated tissue stratification, presenting relevant functional roles and coherent with the findings revealed on the studies from which we have collected the data. It is worth to highlight that, in contrary to the previous studies, MULTILAYER partitions the digital map on the grounds of the contiguous gexel information, it provides to the user ranked lists of over-expressed genes and relevant gene co-expression patterns; thus enhancing the molecular characterization of the spatial information in a selfsupervised manner, similar to those used for tissue image segmentation (reviewed in^19^).

### MULTILAYER allows to process high-resolution ST maps by incorporating a super-Gexel agglomerative compression module

Most of the current available ST maps are issued from glass slides on which barcoded-polyT DNA probes are printed. The manufacturing constraints of these DNA arrays provide a resolution of ~100 μm (equivalent to ~10-40 cells per gexel) with a number of spots ranging between ~1000 to ~5000 (when considering the recent commercial upgrade of the original ST protocol), covering a surface of ~6×6 mm ^3^. An alternative strategy, based on the use of uniquely DNA-barcoded beads deposited onto a glass coverslip enhanced the ST resolution to 10 μm. This methodology known as Slide-Seq, allowed to generate high-resolution ST maps within a circular surface of 3 mm of diameter and ~70 000 uniquely DNA-barcoded beads^20^.

Aiming to use MULTILAYER to analyze high-resolution Slide-Seq maps, but concerned by the technical constraints related to (i) the low number of read counts per gene retrieved within Gexels (**Supplementary Figure 6**) and (ii) the high number of Gexels within the ST map impacting the computational performance (including the display functionalities); we have implemented a complementary script allowing to reduce the ST map complexity. This ad-hoc module, called “MULTILAYER compressor”, generates super-Gexels by agglomerating contiguous gexels defined by a user-provided compression factor. This approach, previously described for image segmentation strategies^21^, allowed to enhance the number of counts per gene within super-Gexels to even comparable levels as those retrieved on regular ST maps (**Supplementary Figure 6**). Furthermore, it reduced the computation performance, such that MULTILAYER could decorticate the functionally relevant digital tissue complexity. This last aspect has been highlighted by the analysis of public Slide-Seq data, including mouse hippocampus and sagittal cortex maps^20,22^. In both cases, raw Slide-Seq maps composed by ~ 70 000 high-resolution gexels, were compressed by factors of 60x, 100x and 175x respectively, leading to grid sizes compatible with the performance of MULTILAYER (**Supplementary Figure 6-9** and **Figure 3**). Normalization and differential gene expression analysis performed on the hippocampus map (reduction factor of 60x), allowed to identify spatially distinct over-expression signatures for factors like PPP3CA (Protein Phosphatase 3 Catalytic Subunit Alpha), the Purkinje Cell Protein 4 (PCP4) or the Synaptosome Associated Protein 25 (SNAP25) (**Supplementary Figure 7 & Figure 3B**), This has been further supported by a global tissue stratification issued from the comparison of gene co-expression patterns retrieved over the whole tissue (43 Spatial communities: **Figure 3C**). The use of higher compression factors (100x and 175x) did not affect the observed over-expression signature for PPP3CA, PCP4 or SNAP25, and increased their related read-counts and differential over-expression levels, in agreement with the agglomerative strategy used for generating super-Gexels (**Supplementary Figure 7C & Figure 3B**). Furthermore, it reduced the number of detected spatial gexel communities (13 and 8 spatial communities for 100x and 175x respectively), but retained the global digital tissue substructures (**Figure 3C, E & G**). Finally, the functional relevance of the various spatial gexel communities were confirmed by Gene Ontology enrichment analysis (ARCHS4 tissue database^23^) assessed for the described three compression factor conditions. (**Figure 3D, F** and **H**).

**Figure 3.**
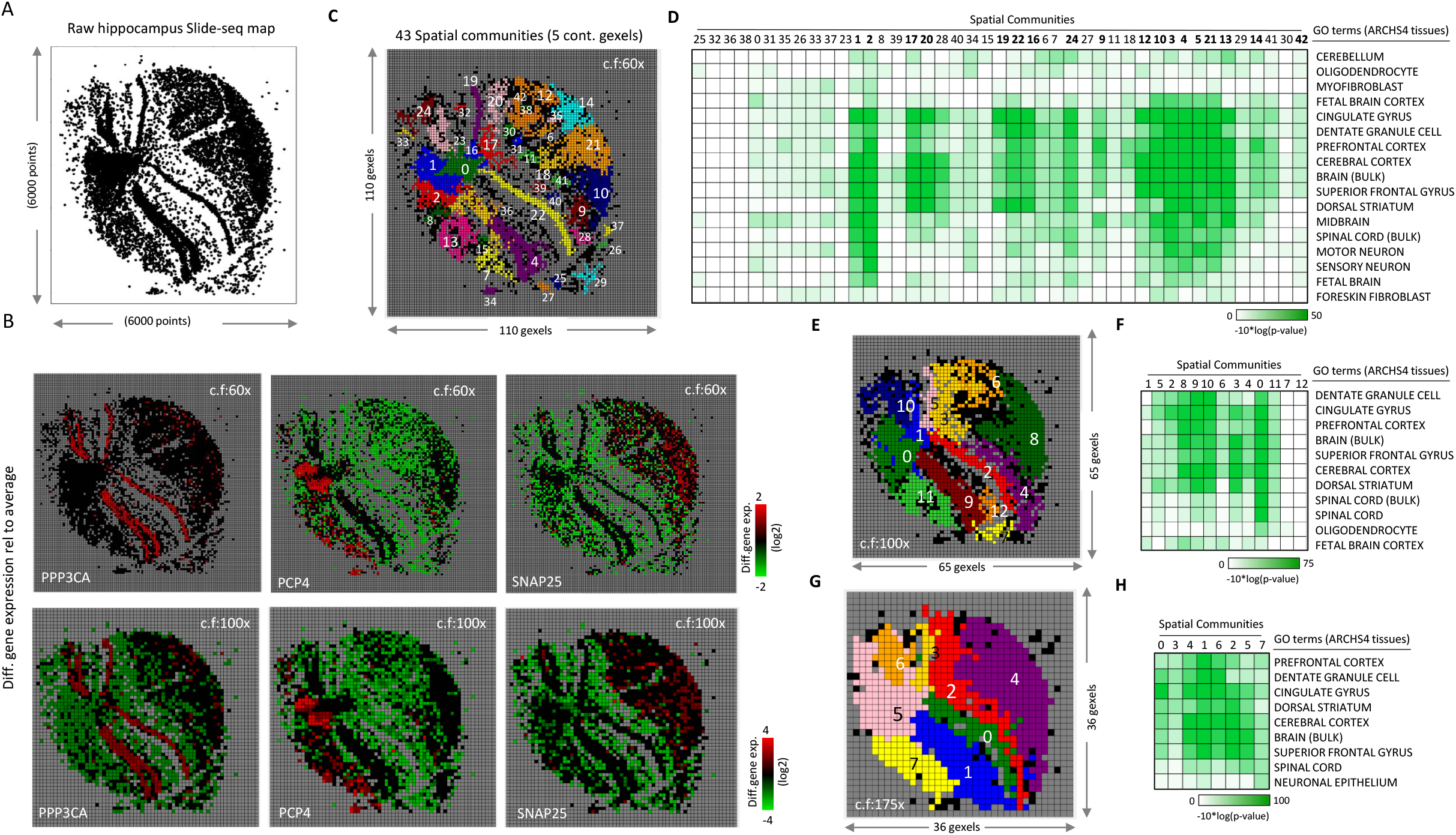
high-resolution hippocampus spatial transcriptomics map analyzed by MULTILAYER. **(A)** Raw hippocampus Slide-seq map displaying the presence of at least one read-count per position. **(B)** Differential expression spatial signatures associated to the factors PPP3CA, PCP4 and SNAP25, after applying a compression factor (c.f.) of 60x and 100x respectively. **(C)** Spatial communities revealed on hippocampus ST map after applying a compression factor of 60x. MULTILAYER compressor reduced the complexity of the original map (displayed in A) to a grid composed by 110×110 super-Gexels which has been processed by MULTILAYER. **(D)** Gene ontology enrichment analysis performed on the 43 spatial communities displayed in (C). **(E & G)** Spatial communities revealed on hippocampus ST map after applying a compression factor (c.f.) of 100x and 175x respectively. **(F & H)** Gene ontology enrichment analysis performed on the spatial communities displayed in (E) & (G) respectively. ARCHS4 tissues: GO terms database issued from Massive Mining of Publicly Available RNA-seq Data from Human and Mouse (*Lachmann A. et al; Nat. Comm. 2018*).

A Similar analysis performed on the cortex map (reduction factor of 60x), allowed to identify spatially distinct over-expression signatures for factors like the gene Transthyretin (TTR) and the Calcium/Calmodulin Dependent Protein Kinase II Inhibitor 1 (CAMK2N1) (**Supplementary Figure 8**), Their gene expression association to the mouse cortex has been confirmed by gene ontology term analysis performed over the spatially co-expressed genes at different digital compression factor reduction levels (100x and 175x in addition to 60x) (**Supplementary Figure 8E-I).** The extension of the co-expression pattern detection over all genes within the tissue map allowed to infer > 40 super-gexel communities on the cortex map issued from a 60x compression factor (**Supplementary Figure 9A**). A gene ontology analysis performed by MULTILAYER (ARCHS4 tissue database) revealed the enrichment for terms like cerebral cortex, superior frontal gyrus, dentate granule cell, motor neuron, neuronal epithelium, dorsal striatum, spinal cord (**Supplementary Figure 9D**). The use of a compression factor of 100x reduced the digital tissue stratification to 15 communities (**Supplementary Figure 9B**) and only to 7 communities when a 175x compression factor has been applied (**Supplementary Figure 9C**). In both cases, the major spatial tissue stratification remained visible, further supported by their associated Gene ontology terms (**Supplementary Figure 9E-F)**.

Overall, MULTILAYER allowed to stratify high-resolution but sparse ST maps, by the use of a super-Gexel agglomerative compression strategy.

### Comparing multiple digitized tissue maps to infer common substructures

A major question to address when counting with multiple tissues issued from related samples is whether their inferred substructures (herein referred as spatial communities) share commonalities enhancing our understanding of their molecular inter-relationship. Recently, Berglund and colleagues have generated ST maps from twelve spatially separated biopsies issued from a cancerous prostate, for which a pathological annotation – based on a histological analysis (Gleason Grading) – has been performed (**Figure 4A**)^24^. To address this question, we have implemented within MULTILAYER a “batch mode”, allowing to process multiple ST maps, but in addition compare all biopsies on the basis of their stratified spatial communities. MULTILAYER inferred spatial community substructures within all 12 sections and revealed their significantly enriched disease-gene associations (**Figure 4B** and **Supplementary Figure 10**).

**Figure 4.**
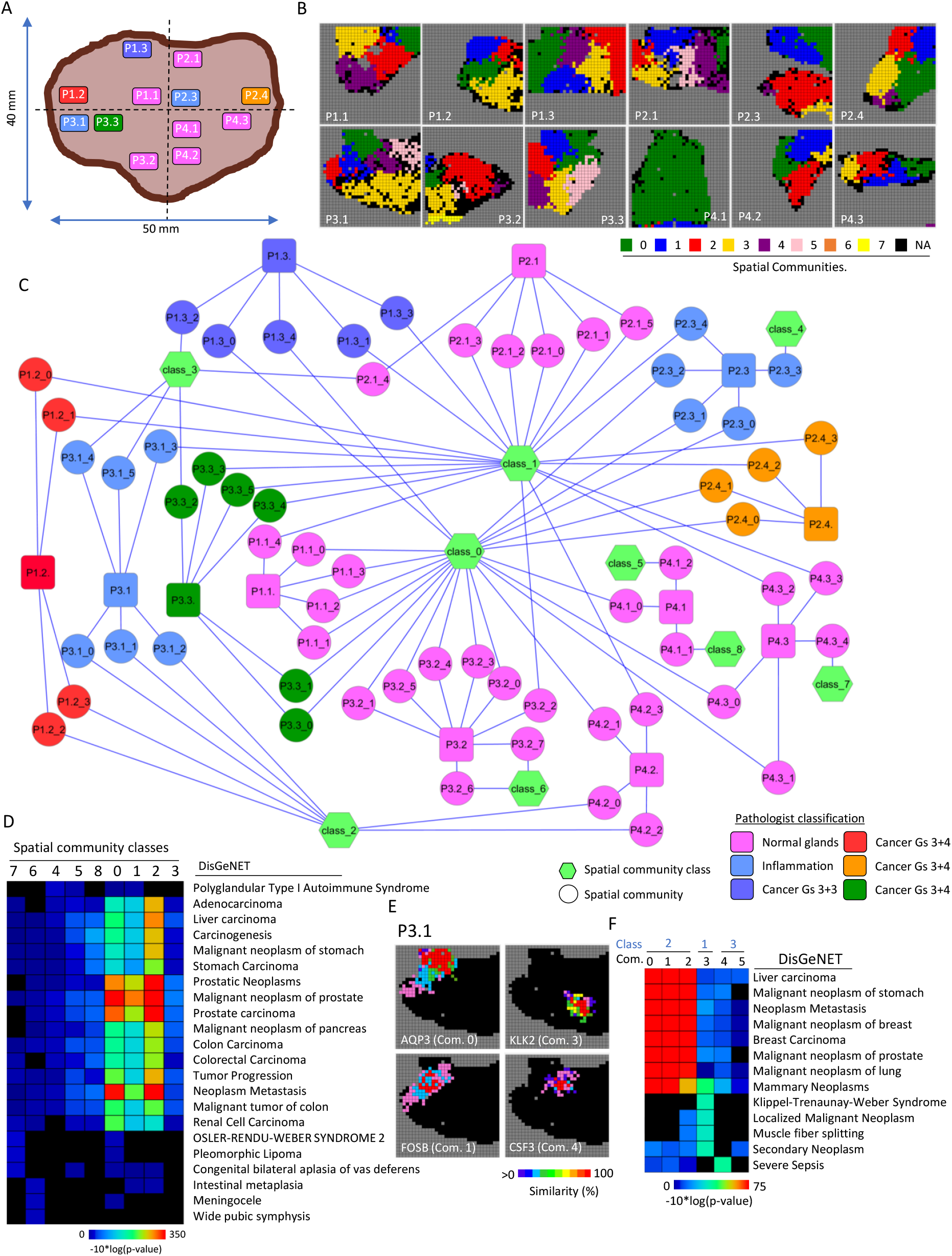
Comparing multiple prostate cancer tissue biopsies to infer common molecular substructures. **(A)** Skim representing the spatial location of 12 tissue biopsies collected from a cancerous prostate and colored in agreement to the histological classification, as described by Berglund et al. **(B)** Spatial transcriptome maps issued from the biopsies illustrated in (A) and processed by MULTILAYER to infer spatial community molecular substructures (color-coded; NA: Non assigned). **(C)** Inter-tissue comparison performed by MULTILAYER (Batch-mode) organizing all spatial communities (round nodes) around 9 “Classes” (green hexagonal nodes). In addition, the tissue biopsy of origin is displayed (rounded-square nodes). Nodes are colored in agreement to the histological classification described by Berglund et al. **(D)** Relevant gene-disease association inferred for the spatial community classes displayed in (C). **(E)** Example of gene co-expression patterns detected on tissue P3.1 at different spatial communities. Gexels in red correspond to the query gene, while others correspond to gene co-expression similarity patterns (Tanimoto index in percent). **(F)** Relevant gene-disease association analysis for each of the spatial communities retrieved on tissue P3.1. The corresponding spatial community classes are also displayed (light blue). DisGeNET: Disease-Gene association discovery platform; Gs: Gleason Score for cancer staging; Com: community.

Inter-tissue comparison was performed by constructing a graph, where spatial communities per tissue are associated to their relevant gene co-expression patterns. Such inter-tissue graph has been partitioned (Louvain methodology^9^) on 9 “classes”, highlighting the relationship between tissue substructures retrieved among all 12 biopsies (**Figure 4C**). Indespite of the histological classification, MULTILAYER partitioning revealed that all tissues present molecular signatures related to prostate cancer progression on at least one substructure (**Supplementary Figure 10, Figure 4C&D**). For instance, tissue biopsies histologically classified as “normal glands” (P1.1, P2.1, P3.2, P4.1, P4.2 and P4.3), presented gene co-expression patterns associated to factors like the membrane cell-junction protein Claudin-4 (CLDN4; known to be over-expressed on primary and metastatic prostate cancer^25^), the growth/differentiation factor-15 (GDF-15; its over-expression has been associated with prostate cancer progression^26^), The gene ACPP coding for the prostatic acid phosphatase, associated to prostatic hyperplasia, but also observed in prostate carcinoma (as revealed on the <u>human protein Atlas database</u>), the Kallikrein Related Peptidase 2 coding for a trypsin-like serine protease (KLK2; primarily expressed in prostate and its overexpression is considered as a prognostic marker for prostate cancer risk^27^), or the activating transcription factor 3 (ATF3), shown to be upregulated on oncogenic stress, and described as a tumor suppressor response notably by its inhibitory effect on androgen receptor signaling^28^ (**Supplementary Figure 10**). Similarly, the tissue P3.1, classified as “inflammation” has been stratified on 6 spatial community substructures, among them 4 being associated to inter-tissue classes functionally enriched to cancer related terms (class 2: communities 0, 1 and 2; class 1: community 3) (DisGeNET^29^ analysis; **Figure 4D & F**). This annotation is supported by the finding of gene co-expression patterns related to factors like the Fos protein FOSB (known to form transcriptionally active heterodimers with the Jun proteins, and reported as being over-expressed on prostate cancer cell lines^30^, but also on prostate cancer biopsies^24^) or the Kallikrein Related Peptidase 2,KLK2 (**Figure 4E**). Furthermore, while the “inflammation” classification has been supported by the local overexpression of the gene Aquaporin-3 (AQP3; community “0”) (**Figure 4E**), the gene coexpression analysis for this factor revealed the presence of other players within the same community, including the Serine Peptidase Inhibitor Kazal Type 1 (SPINK1), previously described as a marker for a molecular subtype of prostate cancer^31^ (**Supplementary Figure 10**). Finally, the spatial communities “4” and “5” appeared devoid of major cancer related terms (which supports their association to class “3”), but still presenting molecular signatures related to prostate cancer incidence, like sever sepsis (**Figure 4F**). Indeed, community “4” present the gene co-expression signature related to the Colony Stimulating Factor 3 (CSF3); known to regulate the generation of infection-protective granulocytes and macrophages^32^, an aspect that is in support of the histological classification of this tissue as “inflammation”.

Finally, the tissue biopsies histologically classified as cancer, did not systematically present all spatial communities related to cancer-related terms. Tissues P1.3 and P3.3 displayed some of their community substructures associated to class “3”, like P3.1 (histologically classified as “inflammation”), and P2.1 (histologically classified as “normal glands”), further supporting the necessity of molecular tissue stratification for better defining tumor progression.

## Discussion

While the use of single-cell transcriptomics for studying the molecular complexity of tissues is gaining in popularity, spatial transcriptomics strategies are anticipated to take-over in the following years, notably with efforts to democratize the access to the required physical supports. In fact, while ST is systematically considered a “non-single cell resolution” assay, in reality all single-cell “omics” approaches converge to aggregate multiple cell readouts into clusters to infer their functional relevance. Similarly, most of the computational algorithms applied on ST maps, process gexels as independent units (i.e. by applying clustering strategies classically used on single-cell “omics” assays), thus under-exploring the available spatial information.

Inspired by the efforts on digital image processing relying on contiguous pixels aggregation, we developed MULTILAYER, a stand-alone package, which considers a ST map as an ensemble of gexels, representing a digital view of the processed tissue. Hence MULTILAYER, “rationalize” the spatial information, by analyzing the presence of contiguous gexels presenting the same gene expression behavior, leading to the stratification of the digital map into molecular tissue substructures.

MULTILAYER provides for the first-time a self-supervised strategy for processing ST maps. It highlights relevant over-expressed gene patterns, which leads to spatial tissue partitioning and infers their functionally relevant Gene ontology associations. Due to its “MULTILAYER” architecture, it provides means to process all types of ST maps, including high-resolution data. Furthermore, it provides means for comparing multiple ST maps, an aspect that is currently limited by the few studies presenting the required data, but is anticipated of extreme interest for instance in the context of developmental or molecular diagnostic studies.

Overall, we anticipate that MULTILAYER corresponds to the first version of algorithms dedicated to process “molecular tissues”, which combined with other strategies like singlecell “omics” might allow to reconstitute digital maps of all organs within the human body, but also contribute to the development of molecular diagnostics strategies to be applied within the future progress on personalized medicine.

## Acknowledgements

This work has been supported by Genopole Thematic Incentive Actions funding (ATIGE-2017) and the institutional bodies CEA, CNRS and Université d’Evry-Val d’Essonne.

## Authors contribution

Conceptualization: J.M. & M.A.M.P.; Methodology J.M., B.M. & M.A.M.P.; Software development: J.M. & B.M.; Scientific evaluation: B.M.C. & M.A.M.P.; Writing, review and editing: J.M. B.M.C. & M.A.M.P; Funding acquisition: M.A.M.P.

## STAR Methods

### Normalization

MULTILAYER corrects for differences on total read-counts within gexels since such variations are considered technical artifacts issued from the sample preparation. For it, quantile normalization methodology (previously described for correcting technical variations within RNA-seq assays^6^) is applied as following: a pseudo-count of 1 is added to all gexels to avoid handling null values. Read-counts per gene within gexels are sorted on the basis of their frequency, then the average read-counts across all ranked gexels is computed based on their raking order (i.e. average within the highest, middle or lower values respectively). Finally, the average read-counts are incorporated instead of the original counts and the readcounts distribution is reorganized as initially. As consequence, when adding all read-counts per gexel after normalization, a constant value is retrieved across all gexels, corresponding to an ideal situation in which all coordinates within the digitized tissue are composed by the same sequencing coverage levels.

### Spatial Differential gene expression

Under the hypothesis that the tissue under study is not homogenous, MULTILAYER performs a differential expression analysis for identifying over/under-expressed genes relative to the global behavior within the tissue. For it, the average of the read counts per gene within the tissue is computed, then the read counts per gene within gexels are expressed relative to the average value (log2). Differentially expressed genes within gexels are defined by a threshold value of two folds (1 or −1 in log2) as a default parameter. As part of the “differential expression” panel, MULTILAYER displays a raking of induced or repressed genes based on the number of gexels within the tissue, allowing to identify most relevant over-expressed genes in an intuitive manner.

### Gene expression pattern detection

Like in digital image processing, MULTILAYER applies an iterative agglomerative strategy(<u>sklearn.cluster.AgglomerativeClustering</u>) over contiguous gexels associated to a given upregulated gene. At the end of the process, gene patterns presenting a user-defined minimal number of contiguous gexels are retained for downstream processing (default threshold: 10 contiguous gexels). The parameters in use within the agglomerative clustering method are: number of clusters (n_clusters): None; affinity: Euclidean; linkage: single; distance threshold (distance_threshold): 1.5.

### Gene co-expression patterns similarity

Having detected gene patterns over the whole tissue, MULTILAYER compares their localization to assess their relevant spatial co-expression. For it, two similarity metrics are implemented on MULTILAYER: Tanimoto/Jaccard and Dice/Sorensen similarity index. Specifically, gene co-expression pattern similarity is evaluated as following:

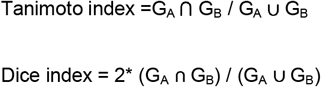

where G_A_ & G_B_ correspond to the number of gexels associated to Gene A and Gene B respectively. All figures presented on this article were obtained using Tanimoto / Jaccard similarity index.

Within the gene co-expression patterns panel of MULTILAYER, over-expressed gene patterns are ranked on the basis of the number of contiguous gexels. Furthermore, the gene co-expression similarity analysis display gexels colored on the basis of their co-expression similarity index, allowing to visualize the extent of co-expressed patterns with the queried gene. In addition, MULTILAYER provides the possibility to perform a Gene Ontology enrichment analysis on the basis of their inferred co-expressed genes (see below).

### Tissue communities’ identification by gene co-expression patterns partitioning

Gene co-expression patterns detected over the whole tissue are represented within MULTILAYER as a major network composed by nodes representing the assessed gene patterns and edges revealing their degree of similarity. This Complex graph is stratified in high modularity community partitions by applying the Louvain hierarchical clustering algorithm^9^. Due to the non-deterministic nature of the Louvain algorithm, MULTILAYER partitions the graph multiple times (15 events by default), then it selects the most frequent community partitions outcome (the frequency of the community partitions are displayed in the terminal), for their display within the tissue map where gexels are colored in agreement to their associations to the inferred communities. In addition, the communities’ panel within MULTILAYER displays the list of over-expressed genes composing the patterns associated to the illustrated communities

Arguments for Louvain partitioning: (i) Weight: allows to include the similarity index computed within the co-expressed genes as a weight argument; (ii) Multiple iterations: Allows to perform 15 consecutive partitioning with the Louvain algorithm and select the most represented for the downstream analyses.

### Gene ontology analysis

MULTILAYER counts with a Gene Ontology enrichment analysis implemented within the “gene co-expression patterns detection” and “Gexel communities” panels. For it, a collection of GO terms has been collected from the <u>Enrichr libraries</u> suite. MULTILAYER infers the GO terms enrichment confidence by comparing the list of genes issued from the gene coexpression patterns detection or within a spatial community, with those retrieved within the GO database (one-sided Fisher Exact test).

As outcome, MULTILAYER provides a confidence barplot per enriched GO terms, as well as a heatmap matrix displaying the list of genes associated to the enriched GO terms.

### MULTILAYER Compressor ad-hoc module

Multilayer Compressor is an ad-hoc module for generating super-Gexel maps by aggregating the raw read counts of contigous gexels prior processing. This strategy allows to convert a larger matrix, like those retrieved in the case of high-resolution Slide-Seq data^20^, to a compressed format, counting with less number of gexels within the grid but with an enhanced number of read-counts per gexel. According to the user-defined compression factor parameters (number of gexels on X & Y coordinates), Multilayer Compressor transforms an input data (3 columns format composed by gexel coordinate, Gene ID and read counts per gene), into a data-frame compatible with MULTILAYER (matrix format composed by gexel coordinates on columns and Gene ID on rows). We recommend to use the MULTILAYER Compressor when raw ST maps are bigger than 120×120 gexel grids.

### Data availability

All processed spatial transcriptomics data within this article were obtained from public repositories and converted into a uniform format compatible with MULTILAYER requirements. Human heart development data generated by Asp. M. et al^10^; and prostate cancer data generated by Berglund et al^24^; were obtained from the <u>Spatial Research portal</u>. Hyppocampus and Cortex high-resolution Slide-seq maps^20^ were obtained from the <u>SpatialDB database</u>^22^. Pancreatic ductal adenocarcinoma data generated by Moncada et al^33^; were obtained from GEO database (<u>GSE111672</u>). Compatible MULTILAYER versions of these data are accessible via https://github.com/SysFate/MULTILAYER.

### Code availability

MULTILAYER and MULTILAYER Compressor are available at https://github.com/SysFate/MULTILAYER.

